# Enemy release mitigates inbreeding depression in native and invasive *Silene latifolia* populations: experimental insight into the role of inbreeding × environment interactions in invasion success

**DOI:** 10.1101/401430

**Authors:** Karin Schrieber, Sabrina Wolf, Catherina Wypior, Diana Höhlig, Stephen R. Keller, Isabell Hensen, Susanne Lachmuth

**Affiliations:** Bielefeld University, Faculty of Biology, Department of Chemical Ecology, 33615 Bielefeld, Germany; Martin-Luther-University Halle-Wittenberg, Institute of Biology, Geobotany & Botanical Garden, 06108 Halle (Saale), Germany; University of Vermont, Department of Plant Biology, Burlington, VT 05405, U.S.A.; German Centre for Integrative Biodiversity Research (iDiv) Halle-Jena-Leipzig, 04103 Leipzig, Germany

**Keywords:** biological invasion, genetic differentiation, genetic paradox, herbivory, purging, white campion

## Abstract

Inbreeding and enemy infestation are common in plants and can synergistically reduce their performance. This inbreeding × environment (I×E) interaction may be of particular importance for the success of plant invasions if introduced populations experience a release from attack by natural enemies relative to their native conspecifics. Using native and invasive plant populations, we investigate whether inbreeding affects infestation damage, whether inbreeding depression in performance is mitigated by enemy release and whether genetic differentiation among native and invasive plants modifies these I×E interactions. We used the plant invader *Silene latifolia* and its natural enemies as a study system. We performed two generations of experimental out- and inbreeding within eight native (European) and eight invasive (North American) *S. latifolia* populations under controlled conditions using field-collected seeds. Subsequently, we exposed the offspring to an enemy exclusion and inclusion treatment in a common garden in the species’ native range to assess the interactive effects of population origin (range), breeding treatment and enemy treatment on infestation damage as well as plant performance. Inbreeding increased flower and leaf infestation damage in plants from both ranges, but had opposing effects on fruit damage in native *versus* invasive plants. Both inbreeding and enemy infestation had negative effects on plant performance, whereby inbreeding depression in fruit number was higher in enemy inclusions than exclusions in plants from both ranges. Moreover, the magnitude of inbreeding depression in fruit number was lower in invasive than native populations. Our results support that inbreeding increases enemy susceptibility of *S. latifolia*, which magnifies inbreeding depression in the presence of enemies. Enemy release in the invaded habitat may thus increase the persistence of inbred founder populations and thereby contribute to successful invasion. Moreover, our findings emphasize that genetic differentiation among native and invasive plants can shape the magnitude and even the direction of inbreeding effects.

## Introduction

Understanding the forces that promote or prevent species range expansions remains a challenging goal in ecology (Barrett, 2015). During invasion of a new range, populations can be simultaneously exposed to increased inbreeding following founder effects (Schrieber & Lachmuth, 2017) and to substantial alterations in the biotic and abiotic environment (Catford, Jansson, & Nilsson, 2009). Inbreeding and environmental change are known to interact in affecting individual fitness (Fox & Reed, 2011), population growth (Reed, Briscoe, & Frankham, 2002) and colonization abilities (Hufbauer, Rutschmann, Serrate, Vermeil de Conchard, & Facon, 2013). Such inbreeding × environment (I×E) interactions are increasingly perceived as potential determinants of species ranges and their dynamics under global change (Colautti, Alexander, Dlugosch, Keller, & Sultan, 2017; Leimu, Vergeer, Angeloni, & Ouborg, 2010; Reed, Fox, Enders, & Kristensen, 2012; Schrieber & Lachmuth, 2017). Nevertheless, empirical studies on the environmental dependency of inbreeding effects in the context of invasions are scarce (Murren & Dudash, 2012; Rosche et al., 2017), despite their potential relevance for the prediction and management of invasive species.

Invaders can preserve high levels of genetic diversity and sufficient outcrossing rates during their expansion due to e.g. mass introductions and genetic admixture (Hufbauer, 2008; Rius & Darling, 2014). However, numerous invasions were evidently accompanied by repeated population bottlenecks during initial introduction and/or colonization at the leading edge of expansion, which resulted in successive genetic depletion and severe inbreeding in phases most crucial for invasion success (reviewed in Schrieber & Lachmuth, 2017). Inbreeding can reduce fitness in the offspring generation (Angeloni, Ouborg, & Leimu, 2011). Such inbreeding depression arises from an increase in genome-wide homozygosity, which enhances the phenotypic expression of deleterious recessive mutations (dominance) and reduces the expression of heterozygote advantage (over-dominance) (Charlesworth & Willis, 2009). Fitness reductions following inbreeding compromise the establishment and growth of colonizing populations (Hufbauer et al., 2013) and are thus assumed to hamper invasions.

The environmental dependency of inbreeding depression provides a hitherto underappreciated explanation for invasion success in the face of increased inbreeding rates. Abiotic and biotic stressors induce changes in gene expression, protein and metabolite synthesis, which maintain physiological homeostasis and, consequently, fitness under unfavorable environmental conditions (Bundy, Davey, & Viant, 2008). Inbreeding can compromise such stress responses *via* dominance and over-dominance effects, which increases the magnitude of inbreeding depression in stressful environments (Fox & Reed, 2011; Kristensen, Pedersen, Vermeulen, & Loeschcke, 2010) while inbreeding depression remains low to absent in benign environments (Enders & Nunney, 2016; Rosche et al., 2017). Plant invasions are often accompanied by a release from environmental stressors such as resource limitation (Blumenthal, 2006), competition (Mitchell et al., 2006) and natural enemies (Keane & Crawley, 2002). Both inbreeding and stress release occur particularly during the early stages of invasion and towards the leading edge of expansion (Dietz & Edwards, 2006; Mitchell et al., 2006). As a consequence, inbreeding depression may be mitigated in small founder populations that experience a stress release in the non-native range, thus fostering invasion success (Schrieber & Lachmuth, 2017).

Natural enemies are especially important to I×E interactions during invasions, since they have the potential to regulate long-term patterns of host plant abundance, population dynamics and distribution (Maron & Crone, 2006). Moreover, there is ample evidence for the dependency of plant inbreeding depression on rates of infestation by natural enemies (Bello-Bedoy & Núñez-Farfán, 2011; Carr & Eubanks, 2002; Hayes, Winsor, & Stephenson, 2004). Studies quantifying inbreeding depression in native and invasive plant populations in the presence *versus* absence of their natural enemies may thus provide insight into the role of I×E interactions in invasion success. Such studies can also yield information on how phenotypic differentiation among host populations impacts the outcome of I×E interactions, which may help to explain reported inconsistency in their effects on plant performance (Fox & Reed, 2011; Sandner & Matthies, 2016; 2017). During invasions, plant species often evolve differences in performance and defense traits (Orians & Ward, 2010; Whitney & Gabler, 2008). This phenotypic divergence can arise either from adaptive responses to changes in the selective regime, for e.g. climate, competition and natural enemies (Agrawal et al., 2015; Colautti & Barrett, 2013; Zhang & Jiang, 2006), or from neutral processes such as genetic drift and founder effects (Keller & Taylor, 2008; Lachmuth, Durka, & Schurr, 2011; Travis et al., 2007). Both adaptive and non-adaptive genetic differentiation likely also alter the genetic architecture underlying inbreeding depression and its dependency on the environment, e.g. through differences in the accumulation and purging of genetic load (Klopfstein, Currat, & Excoffier, 2006; Schrieber & Lachmuth, 2017).

Here, we investigate the combined effects of inbreeding and enemy infestation on the performance of native and invasive populations of the plant species *Silene latifolia* Poir. (Caryophyllaceae). The species is native to Eurasia and has been introduced to North America in the early 19th century. During its invasion of North America, *S. latifolia* experienced events conducive to the expression of I×E-interactions: introduced plants escaped their natural enemies (Wolfe, 2002) and experienced severe population bottlenecks (Keller, Gilbert, Fields, & Taylor, 2012; Taylor & Keller, 2007) as well as high inbreeding levels in founder populations (Fields & Taylor, 2014; Richards, 2000). Moreover, invasive populations evolved differences in enemy susceptibility and performance (Blair & Wolfe, 2004; Wolfe, Elzinga & Biere, 2004; Keller, Sowell, Neimann, Wolfe & Taylor 2009; Schrieber et al. 2017) making *S. latifolia* ideally suited for examining the impact of genetic differentiation on the outcomes of I×E interactions. We conducted experimental inbreeding and outbreeding within native and invasive *S. latifolia* populations, exposed the offspring to the absence and presence of natural enemies, and measured traits related to growth, reproduction and infestation damage to address the following predictions: i) Inbred plants incur higher infestation damage than outbreds. ii) Plant growth and reproduction are lower in inbreds than outbreds (inbreeding depression) and in the presence than absence of natural enemies (stress). iii) The magnitude of inbreeding effects on growth and reproduction is higher in the presence than in the absence of natural enemies (I×E interaction). iv) The aforementioned individual and combined effects of inbreeding and enemy infestation are modified by the distinct evolutionary histories of native and invasive populations.

## Materials and Methods

### Study system

*Silene latifolia* is a short-lived perennial herb mainly distributed across ruderal habitats. The plant is dioecious and produces sexually dimorphic flowers pollinated by insects. Females develop large numbers of capsules containing several hundred seeds, which lack a specific dispersal syndrome and are thus mainly dispersed passively and by human activities. Limited seed dispersal and restricted pollen transfer among neighboring plants can lead to restricted gene flow and the formation of kin-structured patches within populations (McCauley, 1997; 1994). These characteristics have been shown to result in high levels of biparental inbreeding in small, isolated or recently founded *S. latifolia* populations (Fields & Taylor, 2014; Richards, 2000).

In its native range (Eurasia), *S. latifolia* is attacked by three specialist enemies: *Hadena bicruris* Hufn. (Noctuidae) - a noctuid moth that is a specialist pollinator (adult) and a seed predator (larva) at the same time; *Microbotryum violaceum* (Pers.) G. Deml & Oberw. (Mycrobotryaceae) - a systemic sterilizing fungus; and *Brachycaudus lychnidis* L. (Aphididae) - an aphid that causes flowers to abort due to phloem-feeding (Wolfe, 2002). Moreover, native populations are attacked by various leaf- and flower feeding generalist herbivores, including slugs (mainly *Arion lusitanicus* Mabille (Arionidae)), beetles, thrips, caterpillars (often *Mamestra brassicae* L. (Noctuidae)) and leaf miners as well as by several generalist rust and mildew fungi (Schrieber et al. 2017). In the invaded range (North America), *H. bicruris* is completely absent (Wolfe, 2002), the occurrence of *M. violaceum* is locally restricted to a small region in Virginia (Antonovics, Hood, Thrall, Abrams, & Duthie, 2003), and the abundance of aphids as well as leaf and flower feeding generalists is very low relative to the native range (Wolfe, 2002). Invasive *S. latifolia* populations exhibit higher growth and reproduction as well as higher susceptibility to enemy infestation relative to native populations, which can be attributed to both adaptive and non-adaptive evolutionary processes (Blair & Wolfe 2004; Wolfe et al. 2004; Keller et al. 2009; Schrieber et al. 2017). A trade-off between growth/reproduction and enemy susceptibility was not detected in this species (Schrieber et al. 2017).

### Field sampling and experimental setup

We collected open-pollinated seeds from eight native and eight invasive *S. latifolia* populations (Supporting Information Fig. S1, Table S2). Sampling in the native range comprised the geographic source regions of introduction (broadly, eastern and western Europe), while sampling in the invasive range comprised the geographic regions of initial introduction and early expansion (eastern North America), as identified by Taylor & Keller, (2007) and Keller et al. (2012). Within each population, we sampled one capsule (maternal family) from each of five different female plants that were equally distributed over the population area and spatially separated from each other as far as possible. Using these maternal families, we conducted two generations of experimental inbreeding and outbreeding within all native and invasive populations under controlled greenhouse conditions. The offspring were exposed to the absence and presence of natural enemies in a common garden in the species’ native range. Data for the outbred plants from this experiment have previously been used to investigate adaptive and non-adaptive differentiation in growth, reproduction and enemy susceptibility between the native and invaded range (Schrieber et al., 2017).

### Experimental inbreeding and outbreeding

For the P-generation, we germinated ten seeds from each of the five field-collected families in 0.8 mM Giberellic acid in a germination chamber (16 h light at 25 °C, 8 h dark at 13 °C). After six days, the seedlings were planted into pots and transferred to the greenhouse (16 h light at 25 °C, 8 h dark at 13 °C) where they received weekly fertilization (Kamasol Brilliant Rot, Compo Expert, Münster, GE). After seven weeks, we randomly chose one male and one female plant per family for crosses. Each female received pollen from a sib male belonging to the same family (inbreeding), and pollen from a male belonging to a different family within the same population (outbreeding) at distinct flowers (Fig. 1). The crossing of the P-generation resulted in 160 population (N = 16) × family (N = 5) × breeding treatment (N = 2) combinations (PFBCs). For the second generation, we randomly chose one capsule per PFB and propagated the F1-plants from its seeds as described for the P-generation. Female inbred offspring received pollen from an inbred male from the same family, while female outbred offspring received pollen from an outbred male from a different family with respect to the relationships created in the first generation (Fig. 1). We lost seven of the 160 PFBCs due to lack of germination, high mortality, lack of flowering or production of sterile flowers in both inbred and outbred families during the propagation of the F1-generation. Consequently, we obtained a total of 153 PFBCs for the F2-generation plants, which were used for the enemy release experiment.

**Fig. 1:**
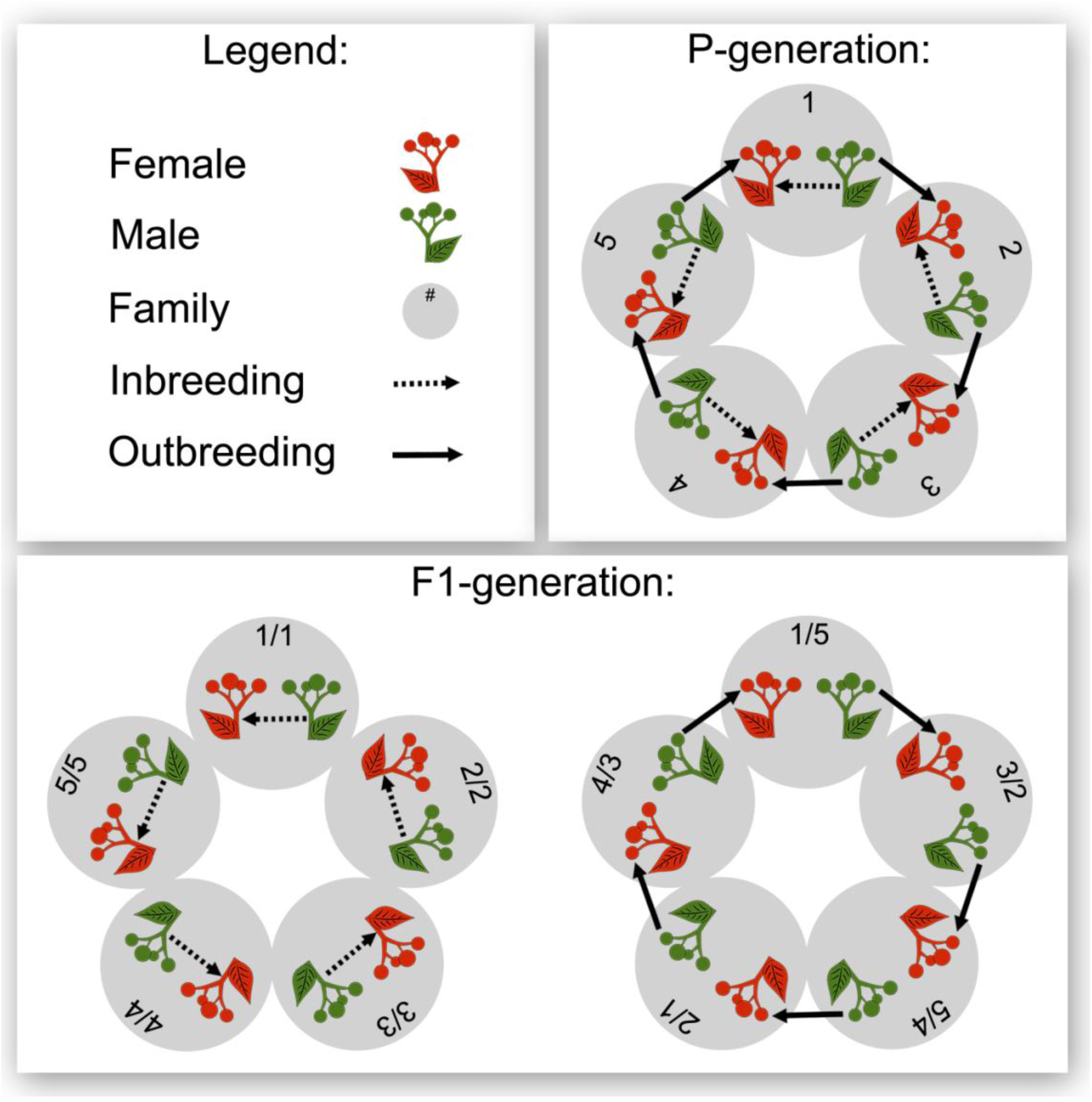
Overview of the two generations of experimental breeding within each of the 16 *Silene latifolia* populations. The crossings were performed with five families (numbered circles). In the P-generation females (orange plants) were fertilized with pollen from males (green plants) from the same family for inbreeding (dashed arrows), and with pollen from males from a different family for outbreeding (solid arrows). In the P-generation, inbreeding and outbreeding were performed at distinct flowers of the same female individual. In the F1-generation, inbreeding was performed with individuals from inbred families and outbreeding with individuals from outbred families from the P-generation. Numbers for the F1-generation families correspond to the maternal/paternal plant of the breedings in the P-generation.

### Enemy release experiment

We exposed native and invasive, inbred and outbred *S. latifolia* plants from the F2-generation to an enemy exclusion and an enemy inclusion treatment using a fully factorial experimental approach (16 populations [8 native *versus* 8 invasive] × 4-5 families × 2 breeding treatments [inbred *versus* outbred] × 2 enemy treatments [exclusion *versus* inclusion] × 8 replicates = 1,224 plants). In early spring, we germinated eight seeds originating from one capsule per PFBC and reared the F2-plants for six weeks in a common garden in Halle (Saale), Germany (51.489 °N 11.959 °E alt: 88 m). After six weeks, we moved the plants to the UFZ Research Station in Bad Lauchstädt, Germany (51.391°N, 11.878°E, alt: 116 m). The planting area was densely covered by a diverse plant community of grasses and forbs including a patchy population of *S. latifolia* that was infested by all of the above-mentioned specialist and generalist enemies. In the common garden, we established four vegetation-free belts, which comprised four plots respectively (∑ = 16 plots) (Fig. 2).

**Fig. 2:**
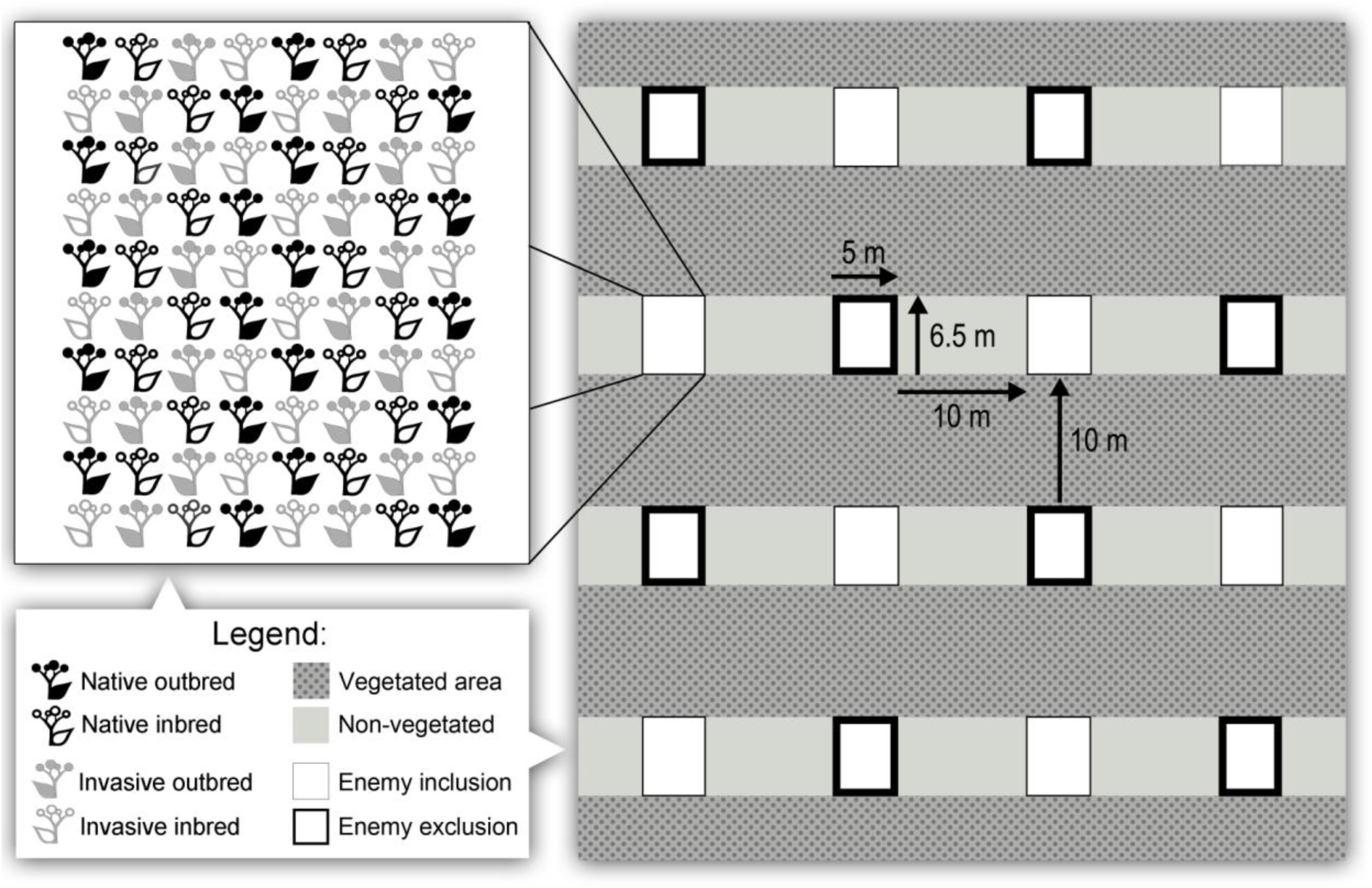
Overview of the experimental manipulation of enemy infestation. The figure illustrates the non-vegetated areas (light gray faces) with the experimental plots (white faces) and the vegetated areas (structured, dark gray faces) from which natural enemies colonized the plots. Either the enemy exclusion (bold black frames) or the enemy inclusion (thin black frames) treatment was applied to each eight uniformly distributed plots. Within each plot, plants were equally distributed with respect to range (native = black, invasive = gray) and breeding treatment (filled = outcrossed, open = inbred).

Each plot included all native and invasive populations represented by two to three F2 maternal families each with one inbred and one outbred individual. As such, the five families within each population were split between two plots (plot pair), which together comprised all of the 153 PFBCs. Each plot pair was replicated an additional seven times. While populations and families were planted randomly within the plots, the range and breeding treatments were uniformly distributed according to a fixed scheme (Fig. 2) in order to reduce confounding plot edge effects. Plots within pairs and plot pair repetitions were randomly distributed across the experimental area. We experimentally excluded natural enemies in eight of the plots (enemy exclusions) over a period of three months (Fig. 2). For this purpose, we used slug fences coated with a gastropod deterrent (Schneckenabwehrpaste, Irka, Mietingen, GE), as well as a molluscicide (Limex, Celaflor), systemic insecticides (alternating between Calypso and Confidor, Bayer, Leverkusen, GE) and a systemic universal fungicide (Baycor M, Bayer, Leverkusen, GE), which were applied in a two-week cycle in accordance with the manufacturers instructions. The remaining eight plots (enemy inclusions) were not treated with pesticides and therefore extensively colonized by specialist and generalist herbivores two weeks after the experiment was set up. The removal of vegetation however deterred *A. lusitanicus* from entering the inclusion plots, so we equipped them with slug fences whose impassable sides were turned towards the plot interior and introduced 15 *A. lusitanicus* individuals to each plot. We adjusted the number of slugs within each inclusion plot to 15 three times a week. The infection with specialist and generalist fungi remained low in all inclusion plots for the entire experimental period. All plots were weeded weekly and watered when necessary during the experiment.

After three months of enemy treatment application, we collected data on morphological defense and infestation damage for each plant in the enemy inclusion plots. We collected leaves at similar stages of development to determine trichome density in a 5 × 5 mm area away from the main vein and at the broadest section of the leaf. In addition, we determined the proportion of flowers (including buds) damaged by tissue removal (generalist herbivores) or phloem sucking (*B. lychnidis)*, the proportion of fruits predated by *H. bicruris* larvae and the proportion of fully grown leaves infested by generalist herbivores (mainly *A. lusitanicus* and *M. brassicae*). Data on infection rates with the specialist fungus *M. violoceaum* and other generalist fungi were not included in the data analysis, as the abundance of these pathogens was generally very low. Furthermore, we collected data on plant growth and reproduction in both enemy inclusion and exclusion plots. We measured the corolla diameter of the biggest flower and counted the number of flowers (including buds) for all male and female plants. Moreover, we determined the number of fruits for all female plants. Seed number, weight and germination were not assessed, since seeds resulted from uncontrolled crossings among plants from all populations, both ranges and both breeding treatments. Finally, we determined the dry aboveground biomass (48 h, 80 °C) for all plant individuals.

### Statistical analysis

All statistical analyses were conducted with R version 3.2.3 (R Development Core Team, 2015). We used linear mixed-effects models (LMMs) for response variables with Gaussian error distribution and generalized linear mixed-effects models (GLMMs) for response variables with Poisson or binomial errors (R-package: lme4; Bates, Maechler, Bolker, & Walker, 2014).

The models for the responses trichome density (LMM, Gaussian, square-root transformed), leaf damage (GLMM, binomial), flower damage (GLMM, binomial) and fruit damage (GLMM, binomial) (all assessed in enemy inclusion plots only) comprised the fixed effects of range and breeding treatment as well as an interaction among both factors. The models for the responses biomass (LMM, Gaussian, square-root transformed), corolla size (LMM, Gaussian), number of flowers (GLMM, Poisson) and number of fruits (GLMM, Poisson) (all assessed in both enemy exclusions and inclusions) comprised the fixed effects of range, breeding treatment and enemy treatment as well as all possible interactions among these factors. All of the described models additionally involved the latitudinal coordinates of the population of origin (centered and scaled) and plant sex (except for fruit damage and number of fruits) as covariates. Moreover, all models included the random effects of plot, population and paternal plant in P-generation nested within population as well as maternal plant in P-generation nested within population.

All models were fitted with a maximum likelihood approach. GLMMs were then tested for under- and overdispersion (R-package: blemco, Korner-Nievergelt et al. 2015). The GLMMs for leaf damage and number of flowers were overdispersed and consequently complemented by an observational level random factor in order to improve the model fit and avoid biased parameter estimates (Harrison, 2014; 2015). After assuring that (G)LMMs exhibit variance homogeneity and normal distribution of residuals by means of visual inspection (Zuur, Ieno, Walker, Saviliev, & Smith, 2009), we applied step-wise backward model selection to obtain the minimal adequate models. Here, we removed fixed effect terms with *p* > 0.05 based on likelihood ratio tests (Venables & Ripley, 2000). For illustration of the interactive effects of range, breeding treatment and enemy treatment on plant performance responses, we extracted least square means with standard errors from the respective full mixed effects models (Lenth, 2016). In contrast to raw data means and their standard errors, these model estimates account for the specific error distribution of the responses, for the effects of covariates as well as for random effects.

## Results

### Interactive effects of range and breeding treatment on morphological plant defense and infestation damage

The density of leaf trichomes was not significantly influenced by range, breeding treatment, the interaction range × breeding treatment or one of the covariates (Table 1, Fig. 3a). The proportion of damaged leaves was significantly related to range and breeding treatment (Table 1). Invasive plants experienced more leaf damage compared to native plants (*p* < 0.05, *χ*^*2*^ = 5.4) and inbred plants from both distribution ranges suffered stronger from leaf infestation compared to outbreds (*p* < 0.001, *χ*^*2*^ = 41.7) (Fig. 3b). The proportion of damaged flowers depended significantly on range, breeding treatment and the covariate sex (Table 1). Flower infestation was higher for invasive than native (*p* < 0.01, *χ*^*2*^ = 6.8), inbred than outbred (*p* < 0.001, *χ*^*2*^ = 41.0) (Fig. 3c) and male than female plants (*p* < 0.05, *χ*^*2*^ = 5.2). The proportion of damaged fruits was significantly influenced by the interaction range × breeding treatment (*p* < 0.05, *χ*^*2*^ = 4.1). Here, invasive plants received generally more fruit damage than native plants and fruit infestation was higher on inbred than outbred native plants but lower on inbred than outbred invasive plants (Fig. 3d).

**Table 1:** Overview and results of analyses evaluating the interactive effects of range, breeding treatment and enemy treatment as well as the effects of covariates (sex, latitudinal origin of population) on the performance of *Silene latifolia*. Responses are presented with their error distribution and the applied model type (LMM, linear mixed effects model; GLMM, generalized linear mixed effects model). The table presents *X*^*2*^ and *p-*values obtained during stepwise backward model selection based on likelihood ratio tests for each fixed effect term as well as the number of groups within random effects and the number of observations for each response; ii: fixed effect term in significant interaction, -: fixed or random effect not tested.

| Responses: | Leaf trichomes |  | Leaf damage |  | Flower damage |  | Fruit damage |  | Biomass |  | Corolla diameter |  | Number flowers |  | Number fruits |  |
| --- | --- | --- | --- | --- | --- | --- | --- | --- | --- | --- | --- | --- | --- | --- | --- | --- |
|  | LMM (Gaussian) |  | GLMM (binomial) |  | GLMM (binomial) |  | GLMM (binomial) |  | LMM (Gaussian) |  | LMM (Gaussian) |  | GLMM (Poisson) |  | GLMM (Poisson) |  |
| Fixed effects: | $\chi^2$ | $p$ | $\chi^2$ | $p$ | $\chi^2$ | $p$ | $\chi^2$ | $p$ | $\chi^2$ | $p$ | $\chi^2$ | $p$ | $\chi^2$ | $p$ | $\chi^2$ | $p$ |
| Range | 0.345 | 0.557 | **5.391** | **0.020** | **6.795** | **0.009** | i.i. | i.i. | i.i. | i.i. | 1.674 | 0.196 | 1.643 | 0.200 | i.i. | i.i. |
| Breeding | 0.213 | 0.644 | **41.693** | **<0.001** | **40.982** | **<0.001** | i.i. | i.i. | **116.629** | **<0.001** | **54.468** | **<0.001** | **24.516** | **<0.001** | i.i. | i.i. |
| Enemy | - | - | - | - | - | - | - | - | i.i. | i.i. | 1.752 | 0.186 | 2.914 | 0.088 | i.i. | i.i. |
| Range × breeding | 0.758 | 0.384 | 0.031 | 0.860 | 1.478 | 0.224 | **4.124** | **0.042** | 3.039 | 0.081 | 0.005 | 0.941 | 0.740 | 0.389 | **5.871** | **0.015** |
| Range × enemy | - | - | - | - | - | - | - | - | **4.772** | **0.029** | 0.191 | 0.663 | 2.651 | 0.103 | 0.301 | 1.070 |
| Breeding × enemy | - | - | - | - | - | - | - | - | 0.058 | 0.810 | 0.091 | 0.763 | 0.340 | 0.557 | **4.150** | **0.042** |
| Range × breeding × enemy | - | - | - | - | - | - | - | - | 0.239 | 0.625 | 0.036 | 0.849 | 0.100 | 0.745 | 0.476 | 0.490 |
| Sex | 0.196 | 0.658 | 0.144 | 0.704 | **5.225** | **0.022** | - | - | **44.511** | **<0.001** | **41.416** | **<0.001** | **133.510** | **<0.001** | - | - |
| Latitudinal origin Population | 1.034 | 0.309 | 1.993 | 0.158 | 1.332 | 0.248 | 2.862 | 0.091 | 0.007 | 0.933 | 0.002 | 0.969 | 0.000 | 0.962 | 1.304 | 0.253 |
| Random effects: | Groups |  | Groups |  | Groups |  | Groups |  | Groups |  | Groups |  | Groups |  | Groups |  |
| Plot | 8 |  | 8 |  | 8 |  | 8 |  | 16 |  | 16 |  | 16 |  | 16 |  |
| Population | 16 |  | 16 |  | 16 |  | 16 |  | 16 |  | 16 |  | 16 |  | 16 |  |
| Population/mother in P-generation | 76 |  | 76 |  | 76 |  | 73 |  | 76 |  | 76 |  | 76 |  | 76 |  |
| Population/father in P-generation | 79 |  | 79 |  | 79 |  | 75 |  | 79 |  | 79 |  | 79 |  | 79 |  |
| Population/observation | - |  | 592 |  | - |  | - |  | - |  | - |  | 1192 |  | - |  |
| Observations | 551 |  | 592 |  | 571 |  | 282 |  | 1192 |  | 1128 |  | 1192 |  | 579 |  |

**Fig. 3:**
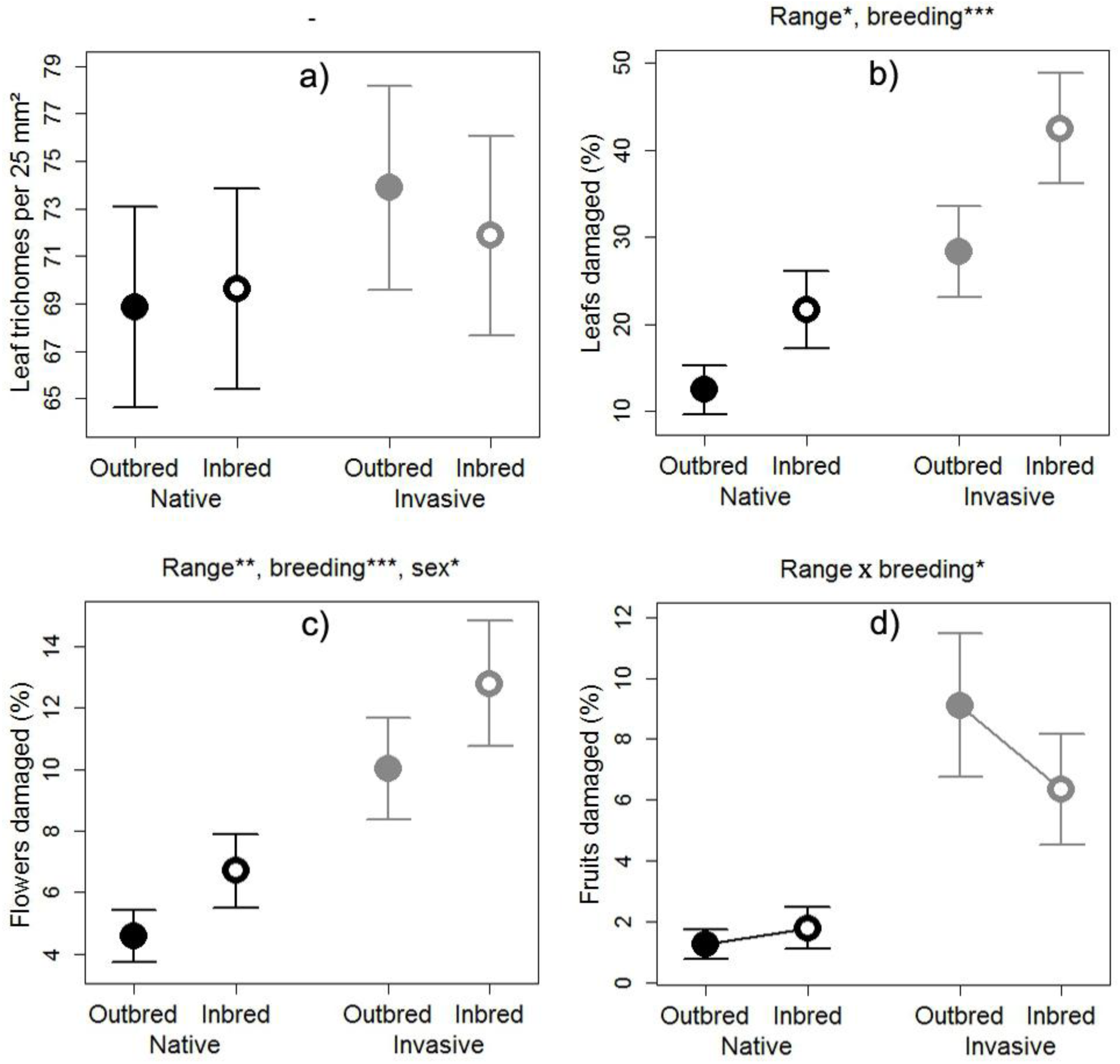
Combined effects of range (native [black] *versus* invasive [gray]) and breeding treatment (outbred [filled] *versus* inbred [open]) on **a)** the density of leaf trichomes, **b)** the proportion of damaged leaves, **c)** the proportion of damaged flowers and **d)** the proportion of damaged fruits in *Silene latifolia*, which were acquired in enemy inclusions. The circles and arrows represent least square means with their standard errors extracted from the full (G)LMMs. The significance levels for fixed effects terms maintained in the minimal adequate (G)LMMs (determined with likelihood ratio tests) are denoted at the top of each plot (*: *p* < 0.05, **: *p* < 0.01, ***: *p* < 0.001). Connecting lines between means of inbreds and outbreds mark significant range × breeding treatment interactions.

### Interactive effects of range, breeding treatment and enemy treatment on plant growth and reproduction

The aboveground biomass of experimental plants was significantly related to the interaction range × enemy treatment, to breeding treatment and to plant sex (Table 1, Fig. 4a). Plants exhibited reduced biomass in enemy inclusions relative to exclusions, whereby this effect was stronger in invasive than native populations (*p* < 0.05, *χ*^*2*^ = 4.8). Inbred plants produced significantly less biomass compared to outbred plants (*p* < 0.001, *χ*^*2*^ = 116.6) and female plants had higher biomass than males (*p* < 0.001, *χ*^*2*^ = 44.5). Range, breeding treatment and enemy treatment had no significant interactive effects on the corolla diameter of *S. latifolia* plants (Table 1). Instead, corolla size was generally lower for inbred than outbred (*p* < 0.001, *χ*^*2*^ = 54.5) (Fig. 4b) and female than male plants (*p* < 0.001, *χ*^*2*^ = 41.4). The number of flowers per plant individual was distinctively lower for inbred than outbred (*p* < 0.001, *χ*^*2*^ = 24.5) (Fig. 4c) and female than male plants (*p* < 0.001, *χ*^*2*^ = 133.5). The number of fruits produced by female plants depended significantly on the two-way interactions range × breeding treatment and breeding treatment × enemy treatment (Table 1, Fig. 4d). Invasive plants produced more fruits than native plants in both breeding and enemy treatments. Moreover, inbred plants had less fruits than outbred plants in both enemy treatments and in populations from both distribution ranges. This inbreeding depression was less intense in invasive than native populations (*p* < 0.05, *χ*^*2*^ = 5.9) and stronger in enemy inclusions than exclusions (*p* < 0.05, *χ*^*2*^ = 4.1).

**Fig. 4:**
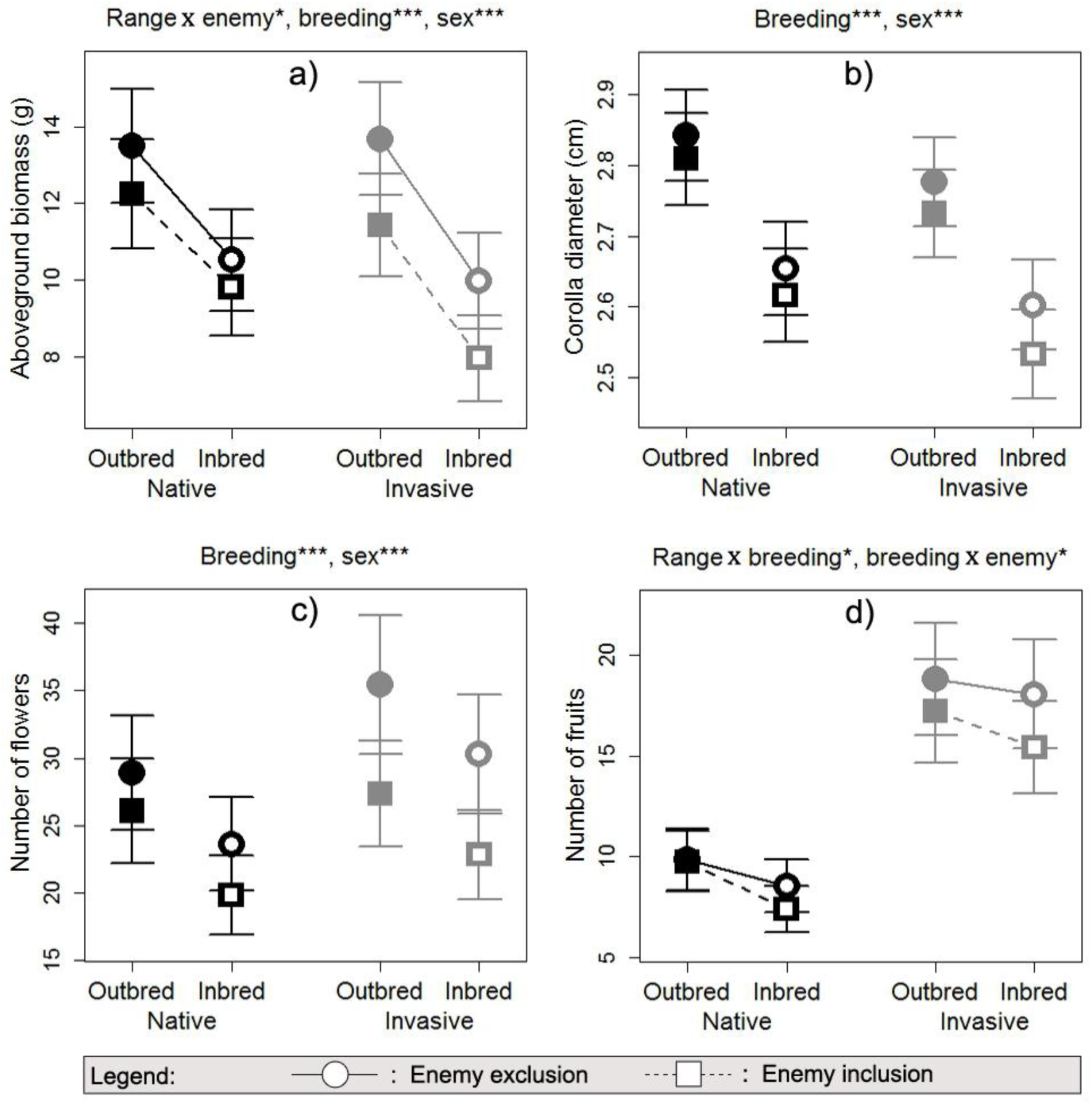
Combined effects of range (native [black] *versus* invasive [gray]), breeding treatment (outbred [filled] *versus* inbred [open]) and enemy treatment (exclusion [circles] *versus* inclusion [squares]) on the **a)** aboveground biomass, **b)** corolla diameter, **c)** number of flowers, and **d)** number of fruits of *Silene latifolia*. The circles and arrows represent least square means with their standard errors extracted from the full (G)LMMs. The significance levels for fixed effects terms maintained in the minimal adequate (G)LMMs (determined with likelihood ratio tests) are denoted at the top of each plot (*: *p* < 0.05, **: *p* < 0.01, ***: *p* < 0.001). Connecting lines between means of inbreds and outbreds mark significant two-way interactions in which breeding treatment is involved.

## Discussion

Our study provides support that I×E interactions can contribute to successful plant invasion and that these interactions are shaped by the evolutionary histories of plant populations. Here, we discuss a) that inbreeding increases enemy infestation damage, which in turn magnifies inbreeding depression in *S. latifolia* plants from both distribution ranges; and b) that some of the inbreeding effects on infestation damage and reproductive traits differ in their magnitude and even in their direction among native and invasive plants as a result of non-adaptive/ adaptive evolutionary processes.

### Enemy release mitigates inbreeding depression in native and invasive *S. latifolia* plants

In accordance with our hypothesis, inbred *S. latifolia* plants from both distribution ranges for the most part incurred higher infestation damage from natural enemies in the common garden than outbreds (Fig. 3b, c; but see Fig. 3d and discussion in next section). Plants often exhibit increased susceptibility to enemies following inbreeding (Bello-Bedoy & Núñez-Farfán, 2011; Campbell, Thaler, & Kessler, 2013; Kariyat, Mena-Alí, et al., 2012), since dominance and over-dominance can either affect gene-loci that contribute directly to plant resistance against enemies (Kariyat, Mena-Alí, et al., 2012) or induce general stress responses that trade-off against responses to environmental stressors such as natural enemies (Kristensen et al., 2010). Using the same inbred and outbred families of native and invasive *S. latifolia* populations investigated in the present study, Schrieber, Kröner, Schweiger and Müller (in press) demonstrated that inbreeding significantly compromises the plants’ chemical responses to insect herbivory. That study also indicated that higher infestation damage on inbred *S. latifolia* individuals can result from compensatory feeding triggered by poor host plant quality. Previous studies on other plant species also demonstrated that inbreeding reduces the concentration of chemicals mediating direct (Campbell et al., 2013) and indirect (Kariyat, Mauck, De Moraes, Stephenson, & Mescher, 2012) plant defense as well as host plant quality (Leimu, Kloss, & Fischer, 2008).

In line with our expectation, both inbreeding and enemy infestation reduced the growth and reproduction of *S. latifolia* in native and invasive populations, whereby inbreeding had a pronounced impact while the effect of enemy infestation was more moderate (Fig. 4). As hypothesized, the effects of breeding and enemy treatment were not purely additive. While the magnitude of inbreeding depression was independent of the enemy treatment for biomass, corolla diameter and flower number (Fig 4a, b, c), it was significantly lower in enemy exclusions than inclusions for fruit number in both distribution ranges (Fig. 4d). The observation that inbreeding depression in traits closely linked to individual fitness increases under stress, while morphological traits that are only indirectly related to reproductive success (i.e., biomass, corolla diameter) are not significantly affected by I×E interactions, has also been made in previous studies (Bello-Bedoy & Núñez-Farfán, 2011; Schou, Loeschcke, & Kristensen, 2015). This difference can indeed be expected, since the investment in reproduction by the end of a growing season is highly dependent on an individual’s cumulative performance and thus cumulative (interactive) effects of inbreeding and herbivory on performance throughout the season (Orr, 2009). Moreover, in contrast to flower number, fruit number in experimental *S. latifolia* plants was affected by all three types of infestation damage (leaf, flower and fruit infestation) and thus pressure by natural enemies was highest for this fitness related trait. Our finding indicates that the enemy release in the invaded habitat may mitigate detrimental inbreeding effects on reproductive output in colonizing *S. latifolia* populations. The I×E interactions detected for native populations under experimental conditions may be representative of a scenario of initial population founding and establishment during early invasion phases, in which plants are naive to the novel environment. I×E interactions in the investigated invasive plants, in turn, may represent a scenario of population founding at the leading edge, where populations have already undergone evolutionary responses to the environment of the introduced range.

Our findings of lower inbreeding depression in benign *versus* harsh habitats in a plant invader go along with those of two previous studies on *Mimulus guttatus* DC. (Murren & Dudash, 2012) and *Centaurea stoebe* L. (Rosche et al., 2017), which together emphasize the relevance of I×E interactions for species expansions and the need for further research under more realistic field conditions. The transplantation of inbred and outbred plants to native as well as invasive field habitats is necessary to assess the net effect of multiple stressors occurring in both environments on the magnitude of inbreeding depression. Ideally, such approaches should involve the quantification of demographic rates in order to parameterize models that estimate population growth and/or spread rates (Normand, Zimmermann, Schurr, & Lischke, 2014; Schultz, Eckberg, Berg, Louda, & Miller, 2017). Studies of this kind could further elaborate whether and to what extent I×E interactions add to several other mechanisms (e.g., genetic admixture, mass introductions, Estoup et al., 2016; Roman & Darling, 2007) that can explain the successful spread of invaders in the face of genetic bottlenecks, i.e. the so-called genetic paradox of biological invasions (Schrieber & Lachmuth, 2017).

### Evolutionary history modifies the magnitude and direction of inbreeding effects on plant interactions with natural enemies

We detected evolutionary differentiation in plant susceptibility to enemy infestation (Fig. 3b, c,d) and plant performance among native and invasive populations of *S. latifolia* (Fig. 4d). This observation has also been made in previous studies of *S. latifolia* (Blair & Wolfe, 2004; Keller et al., 2009; Schrieber et al., 2017; Wolfe et al., 2004), where it has been discussed in detail with regard to the driving evolutionary forces (i.e. adaptive *versus* non-adaptive evolution, potential selective agents, trade-offs).

However, new to our study is the finding that the magnitude and even the direction of inbreeding effects on some metrics of infestation damage and reproductive traits differed among native and invasive *S. latifolia* populations. While inbreeding slightly increased fruit damage in native plants, the proportion of fruits infested by *H. bicruris* was considerably lower on inbreds than outbreds for invasive plants. At the same time, fruit infestation was generally higher on invasive plants (Fig. 3d). This finding may be attributed to the combined effects of genetic differentiation and inbreeding on host plant attractivity. Previous studies elaborated that higher fruit infestation by *H. bicruris* on invasive *S. latifolia* does not result from higher larva performance, but from increased oviposition rates (Elzinga & Bernasconi, 2009). Adult *H. bicruris* females are attracted to oviposition on *S. latifolia* by a specific floral volatile blend, whereby invasive plants emit larger total amounts of these volatiles (Dötterl, Jürgens, Wolfe, & Biere, 2009). Based on studies using other plant species (Delphia, Rohr, Stephenson, De Moraes, & Mescher, 2009; Ferrari, Stephenson, Mescher, & Moraes, 2006), one possibility is that inbreeding reduced the total floral volatile production in *S. latifolia* either directly by impairing the synthesis of volatiles or by reducing flower number (Fig 4c, enemy exclusions), which may have made inbred plants less attractive hosts for oviposition. This potential inbreeding effect on host plant attractivity may have been only apparent in invasive populations due to a specific volatile concentration threshold for the plant’s apparency to *H. bicruris*. Further studies on the inbreeding effects on the composition and concentration of floral volatiles are necessary to test this assumption.

In addition, we found that inbreeding depression for fruit number was less pronounced in introduced relative to native populations (Fig. 4d). Differences in the magnitude of experimental inbreeding depression among populations have often been related to the history of inbreeding within natural source populations (Angeloni et al., 2011). Inbreeding depression is assumed to be lower in populations that have experienced high levels of natural inbreeding, as a result of exposing deleterious recessive mutations to negative selection. As a consequence, the frequency of these mutations within populations can rapidly decrease (i.e. purging of genetic load; Crnokrak & Barrett, 2002), which reduces dominance effects resulting from experimental inbreeding. Invasive populations of *S. latifolia* indeed experienced increased inbreeding levels during the colonization of North America, as evinced by inter- and intra population crossing experiments (Richards, 2000), enhanced genetic structure in recently founded compared to longer established populations (McCauley, Raveill, & Antonovics, 1995) and the occurrence of severe demographic bottlenecks during initial founding (Keller et al., 2012; Taylor & Keller, 2007). Since reproductive traits such as fruit production are crucial for invasion success (Burns, Ashman, Steets, Harmon-Threatt, & Knight, 2011; Phillips, Brown, & Shine, 2010), strong negative selection against genetic load may have induced highly efficient purging under demographic conditions of colonization and invasion.

## Conclusions

Our findings indicate that stress release during invasions may mitigate inbreeding depression in founding populations, which potentially contributes to the successful establishment and expansion of introduced populations. On the other hand, I×E interactions may hamper the colonization of novel habitats that exhibit increased stress levels relative to a species’ native source habitat (Hufbauer et al., 2013). Furthermore, our data illustrate that the inbreeding effects on an organism’s interaction with its environment are likely shaped by the evolutionary histories of populations. As the native and invaded range of a species can differ systematically in the stress regimes they experience, ongoing invasions provide ideal study systems for investigating the effects of evolutionary differentiation on the outcomes of I×E interactions, and how, in turn, the different outcomes may alter the evolutionary trajectories of invasive populations. Studies addressing these issues hold implications that extend far beyond invasive model species. I×E interactions may potentially shape the dynamics of natural populations whenever they are simultaneously exposed to habitat change and increased inbreeding rates following founder effects or population size reductions. These conditions occur not only during species range expansions, but also during range shifts and retractions in the course of global change (Colautti et al., 2017).

## Acknowledgements

This study was financially supported by the German Academic Exchange Service (DAAD, 315-kp-hus) and the graduate program for the promotion of network building among female Ph.D. students from the Martin Luther University of Halle Wittenberg. We thank Tabea and Chris Steinbeißer-Fitz for accommodation during the fieldwork in the USA. We are grateful for the technical assistance provided by Roman Adler, Eva Bremer, Fréderic Couderc, Sandrine Maurice, Ines Merbach, Tina Klemme, Philipp Kuttig, Birgit Müller, Antonella Vinei, Christine Voigt and Eckhard Winter in field, lab and greenhouse. We also thank Helge Bruelheide and the Helmholtz Centre for Environmental Research (UfZ) Halle for providing space and material for the experiment.

## Author contributions

SL and KS conceptualized the research and designed the study. KS conducted the field sampling, the experimental crossings and the enemy release experiment and collected the data with assistance of CW, DH and SW. KS analyzed the data. KS and SL interpreted the data. KS wrote the first version of the manuscript and SL, SK and IH contributed to the final version.

## Supporting Information

### Supporting Information 1, Fig. S1

Map of the geographic locations of the sampled native and invasive *Silene latifolia* populations.

### Supporting Information 2, Table S2

Overview of the geographic locations and sizes of the sampled native and invasive *Silene latifolia* populations.

### Supporting Information 1

**Fig. S1:**
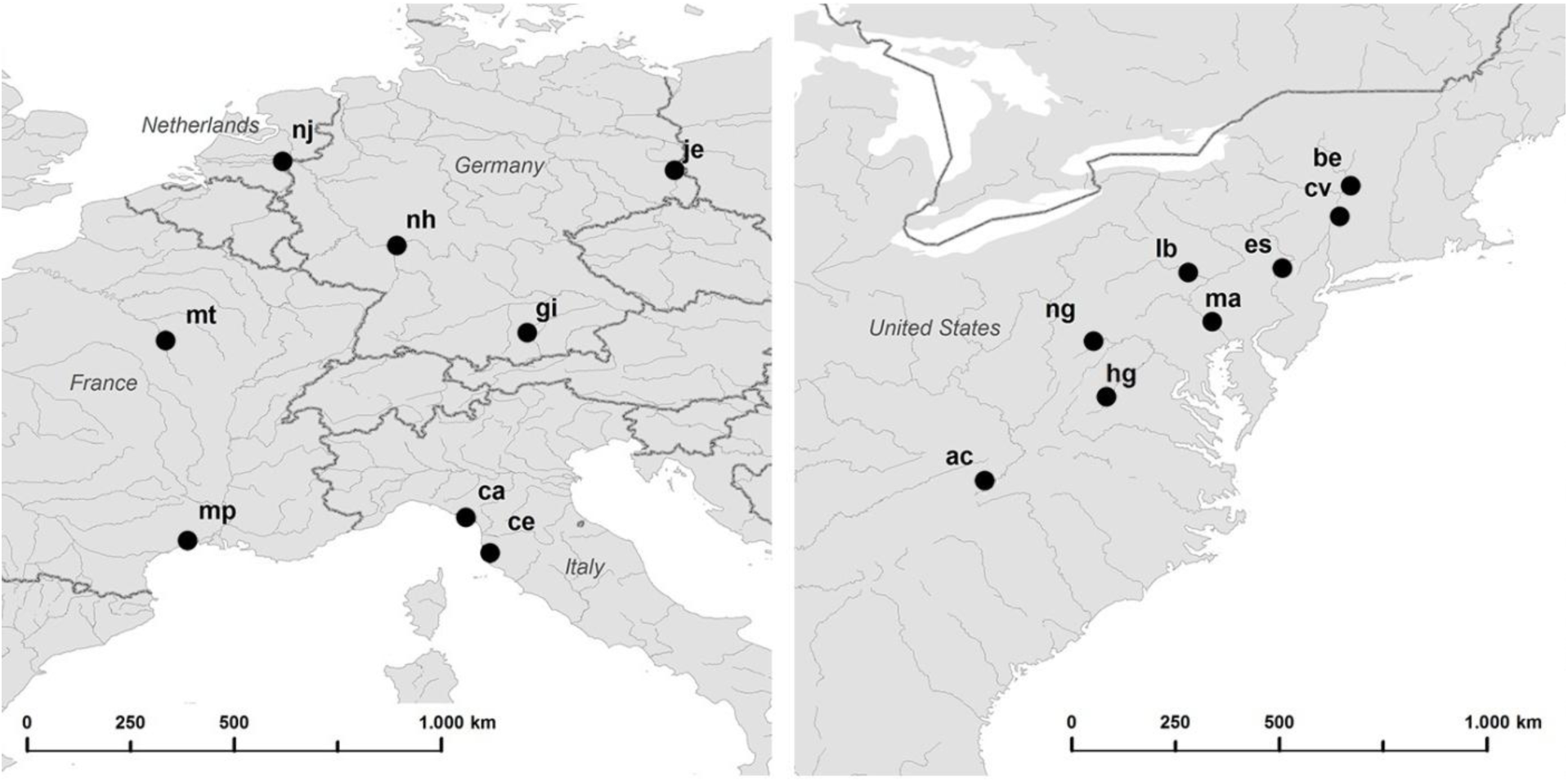
Map of the geographic locations of the sampled native (left) and invasive (right) *Silene latifolia* populations.

### Supporting Information 2

**Table S2:** Overview of the geographic locations and sizes of the sampled native and invasive *Silene latifolia* populations.

| Range | City (state) | ID | °N | °E | Population size |
| --- | --- | --- | --- | --- | --- |
| Native | Caresana (IT) | ca | 44.07702 | 9.96384 | 262 |
| Native | Cecina (IT) | ce | 43.313769 | 10.51195 | 149 |
| Native | Gilching (GE) | gi | 48.105634 | 11.253771 | 35 |
| Native | Jethe (GE) | je | 51.6899 | 14.59683 | 485 |
| Native | Montpellier (FR) | mp | 43.653567 | 3.893247 | 341 |
| Native | Monteneau (FR) | mt | 47.848117 | 3.552508 | 19 |
| Native | Nackenheim (GE) | nh | 49.925119 | 8.342758 | 132 |
| Native | Nijmegen (GE) | nj | 51.883074 | 5.85185 | 46 |
| Invasive | Crumpler (NC) | ac | 36.52576 | -81.41558 | 60 |
| Invasive | Bennington (VT) | be | 42.894717 | -73.294919 | 31 |
| Invasive | Hillsdale (NY) | cv | 42.234825 | -73.506433 | 350 |
| Invasive | Bushkill (PA) | es | 41.14009 | -74.9294 | 900 |
| Invasive | Harrisonburg (VA) | hg | 38.49079 | -78.97762 | 70 |
| Invasive | Lewisburg (PA) | lb | 40.98179 | -76.93041 | 184 |
| Invasive | Washington Boro (PA) | ma | 39.99635 | -76.47243 | 1100 |
| Invasive | Grantsville (MD) | ng | 39.637308 | -79.099481 | 1000 |

## References

Agrawal, A. A., Hastings, A. P., Bradburd, G. S., Woods, E. C., Züst, T., Harvey, J. A., & Bukovinszky, T. (2015). Evolution of plant growth and defense in a continental introduction. The American Naturalist, 186(1), E1–E15. doi:http://doi.org/10.1086/681622.

Angeloni, F., Ouborg, N. J., & Leimu, R. (2011). Meta-analysis on the association of population size and life history with inbreeding depression in plants. Biological Conservation, 144(1), 35–43. doi:10.1016/j.biocon.2010.08.016.

Antonovics, J., Hood, M. E., Thrall, P. H., Abrams, J. Y., & Duthie, G. M. (2003). Herbarium studies on the distribution of anther-smut fungus (Microbotryum violaceum) and Silene species (Caryophyllaceae) in the eastern United States. American Journal of Botany, 90(10), 1522–1531. doi:10.3732/ajb.90.10.1522.

Barrett, S. C. H. (2015). Foundations of invasion genetics: the Baker and Stebbins legacy. Molecular Ecology, 24(9), 1927–1941. doi:10.1111/mec.13014.

Bates, D., Maechler, M., Bolker, B., & Walker, S. (2014). lme4: Linear mixed-effects models using Eigen and S4. R package version 1.1-7. Retrieved from URL: http://CRAN.R-project.org/package=lme4 (2014).

Bello-Bedoy, R., & Núñez-Farfán, J. (2011). The effect of inbreeding on defence against multiple enemies in Datura stramonium. Journal of Evolutionary Biology, 24(3), 518–530. doi:10.1111/j.1420-9101.2010.02185..

Blair, A. C., & Wolfe, L. M. (2004). The evolution of an invasive plant: an experimental study with Silene latifolia. Ecology, 85(11), 3035–3042. doi:10.1890/04-034.

Blumenthal, D. M. (2006). Interactions between resource availability and enemy release in plant invasion. Ecology Letters, 9(7), 887–895. doi:10.1111/j.1461- 0248.2006.00934..

Bundy, J. G., Davey, M. P., & Viant, M. R. (2008). Environmental metabolomics: a critical review and future perspectives. Metabolomics, 5(1), 3–21. doi:10.1007/s11306-008- 0152-.

Burns, J. H., Ashman, T.-L., Steets, J. A., Harmon-Threatt, A., & Knight, T. M. (2011). A phylogenetically controlled analysis of the roles of reproductive traits in plant invasions. Oecologia, 166(4), 1009–1017. doi:10.1007/s00442-011-1929-.

Campbell, S. A., Thaler, J. S., & Kessler, A. (2013). Plant chemistry underlies herbivore-mediated inbreeding depression in nature. Ecology Letters, 16(2), 252–260. doi:10.1111/ele.1203.

Carr, D. E., & Eubanks, M. D. (2002). Inbreeding alters resistance to insect herbivory and host plant quality in Mimulus guttatus (Scrophulariaceae). Evolution, 56(1), 22–30. doi:10.1554/0014-3820(2002)056[0022:IARTIH]2.0.C.

Catford, J. A., Jansson, R., & Nilsson, C. (2009). Reducing redundancy in invasion ecology by integrating hypotheses into a single theoretical framework. Diversity and Distributions, 15(1), 22–40. doi:10.1111/j.1472-4642.2008.00521..

Charlesworth, D., & Willis, J. H. (2009). The genetics of inbreeding depression. Nature Reviews Genetics, 10(11), 783–796. doi:10.1038/nrg266.

Colautti, R. I., Alexander, J. M., Dlugosch, K. M., Keller, S. R., & Sultan, S. E. (2017). Invasions and extinctions through the looking glass of evolutionary ecology. Philosophical Transaction of the Royal Society B: Biological Sciences, 372(1712), 20160031. doi:10.1098/rstb.2016.003.

Colautti, R. I., & Barrett, S. C. H. (2013). Rapid adaptation to climate facilitates range expansion of an invasive plant. Science, 342(6156), 364–366. doi:10.1126/science.124212.

Crnokrak, P., & Barrett, S. C. (2002). Perspective: purging the genetic load: a review of the experimental evidence. Evolution, 56(12), 2347–2358. doi:10.1111/j.0014- 3820.2002.tb00160..

Delphia, C. M., Rohr, J. R., Stephenson, A. G., De Moraes, C. M., & Mescher, M. C. (2009). Effects of genetic variation and inbreeding on volatile production in a field population of horsenettle. International Journal of Plant Sciences, 170(1), 12–20. doi:10.1086/59303.

Dietz, H., & Edwards, P. J. (2006). Recognition that causal processes change during plant invasion helps explain conflicts in evidence. Ecology, 87(6), 1359–1367. doi:10.1890/0012-9658(2006)87[1359:RTCPCD]2.0.CO;.

Dötterl, S., Jürgens, A., Wolfe, L., & Biere, A. (2009). Disease status and population origin effects on floral scent: potential consequences for oviposition and fruit predation in a complex interaction between a plant, fungus, and noctuid moth. Journal of Chemical Ecology, 35(3), 307–319. doi:10.1007/s10886-009-9601-.

Elzinga, J. A., & Bernasconi, G. (2009). Enhanced frugivory on invasive Silene latifolia in its native range due to increased oviposition. Journal of Ecology, 97(5), 1010–1019. doi:10.1111/j.1365-2745.2009.01534..

Enders, L. S., & Nunney, L. (2016). Reduction in the cumulative effect of stress-induced inbreeding depression due to intragenerational purging in Drosophila melanogaster. Heredity, 116(3), 304–313. doi:10.1038/hdy.2015.10.

Estoup, A., Ravigné, V., Hufbauer, R., Vitalis, R., Gautier, M., & Facon, B. (2016). Is There a Genetic Paradox of Biological Invasion? Annual Review of Ecology, Evolution, and Systematics, 47(1), 51–72. doi:10.1146/annurev-ecolsys-121415-03211.

Ferrari, M. J., Stephenson, A. G., Mescher, M. C., & Moraes, C. M. D. (2006). Inbreeding effects on blossom volatiles in Cucurbita pepo subsp. texana (Cucurbitaceae). American Journal of Botany, 93(12), 1768–1774. doi:10.3732/ajb.93.12.176.

Fields, P. D., & Taylor, D. R. (2014). Determinants of genetic structure in a nonequilibrium metapopulation of the plant Silene latifolia. PLOS ONE, 9(9), e104575. doi:10.1371/journal.pone.010457.

Fox, C. W., & Reed, D. H. (2011). Inbreeding depression increases with environmental stress: an experimental study and meta-analysis. Evolution, 65(1), 246–258. doi:10.1111/j.1558-5646.2010.01108..

Harrison, X. A. (2014). Using observation-level random effects to model overdispersion in count data in ecology and evolution. PeerJ, 2, e616. doi:10.7717/peerj.61.

Harrison, X. A. (2015). A comparison of observation-level random effect and Beta-Binomial models for modelling overdispersion in binomial data in ecology & evolution. PeerJ, 3, e1114. doi:10.7717/peerj.111.

Hayes, C. N., Winsor, J. A., & Stephenson, A. G. (2004). Inbreeding influences herbivory in Cucurbita pepo ssp. texana (Cucurbitaceae). Oecologia, 140(4), 601–608. doi:10.1007/s00442-004-1623-.

Hufbauer, R. A., Rutschmann, A., Serrate, B., Vermeil de Conchard, H., & Facon, B. (2013). Role of propagule pressure in colonization success: disentangling the relative importance of demographic, genetic and habitat effects. Journal of Evolutionary Biology, 26(8), 1691–1699. doi:10.1111/jeb.1216.

Hufbauer, R. A. (2008). Biological invasions: paradox lost and paradise gained. Current Biology, 18(6), R246–R247. doi:10.1016/j.cub.2008.01.03.

Kariyat, R. R., Mauck, K. E., De Moraes, C. M., Stephenson, A. G., & Mescher, M. C. (2012). Inbreeding alters volatile signalling phenotypes and influences tri-trophic interactions in horsenettle (Solanum carolinense L.). Ecology Letters, 15(4), 301–309. doi:10.1111/j.1461-0248.2011.01738..

Kariyat, R. R., Mena-Alí, J., Forry, B., Mescher, M. C., Moraes, C. M., & Stephenson, A. G. (2012). Inbreeding, herbivory, and the transcriptome of Solanum carolinense. Entomologia Experimentalis et Applicata, 144(1), 134–144. doi:10.1111/j.1570- 7458.2012.01269..

Keane, R. M., & Crawley, M. J. (2002). Exotic plant invasions and the enemy release hypothesis. Trends in Ecology & Evolution, 17(4), 164–170. doi:10.1016/S0169-5347(02)02499-.

Keller, S. R., Gilbert, K. J., Fields, P. D., & Taylor, D. R. (2012). Bayesian inference of a complex invasion history revealed by nuclear and chloroplast genetic diversity in the colonizing plant Silene latifolia. Molecular Ecology, 21(19), 4721–4734. doi:10.1111/j.1365-294X.2012.05751..

Keller, S. R., Sowell, D. R., Neiman, M., Wolfe, L. M., & Taylor, D. R. (2009). Adaptation and colonization history affect the evolution of clines in two introduced species. New Phytologist, 183(3), 678–690. doi:10.1111/j.1469-8137.2009.02892..

Keller, S. R., & Taylor, D. R. (2008). History, chance and adaptation during biological invasion: separating stochastic phenotypic evolution from response to selection. Ecology Letters, 11(8), 852–866. doi:10.1111/j.1461-0248.2008.01188..

Klopfstein, S., Currat, M., & Excoffier, L. (2006). The fate of mutations surfing on the wave of a range expansion. Molecular Biology and Evolution, 23(3), 482–490. doi:10.1093/molbev/msj05.

Korner-Nievergelt, F., Roth, T., Felten, S., Guelat, J., Almasi, B., & Korner-Nievergelt, P. (2015). Bayesian data analysis in ecology using linear models with R. Elsevier. Retrieved from https://www.R-project.org/package=blmeco

Kristensen, T. N., Pedersen, K. S., Vermeulen, C. J., & Loeschcke, V. (2010). Research on inbreeding in the ‘omic’ era. Trends in Ecology & Evolution, 25(1), 44–52. doi:10.1016/j.tree.2009.06.01.

Lachmuth, S., Durka, W., & Schurr, F. M. (2011). Differentiation of reproductive and competitive ability in the invaded range of Senecio inaequidens: the role of genetic Allee effects, adaptive and nonadaptive evolution. New Phytologist, 192(2), 529–541. doi:10.1111/j.1469-8137.2011.03808..

Leimu, R., Kloss, L., & Fischer, M. (2008). Effects of experimental inbreeding on herbivore resistance and plant fitness: the role of history of inbreeding, herbivory and abiotic factors. Ecology Letters, 11(10), 1101–1110. doi:10.1111/j.1461-0248.2008.01222..

Leimu, R., Vergeer, P., Angeloni, F., & Ouborg, N. J. (2010). Habitat fragmentation, climate change, and inbreeding in plants. Annals of the New York Academy of Sciences, 1195(1), 84–98. doi:10.1111/j.1749-6632.2010.05450..

Lenth, R. V. (2016). Leastsquares means: the R package lsmeans. Journal of Statistical Software, 69(1), 1–33. doi:10.18637/jss.v069.i0.

Maron, J. L., & Crone, E. (2006). Herbivory: effects on plant abundance, distribution and population growth. Proceedings of the Royal Society of London B: Biological Sciences, 273(1601), 2575–2584. doi:10.1098/rspb.2006.358.

McCauley, D. E. (1997). The relative contributions of seed and pollen movement to the local genetic structure of Silene alba. Journal of Heredity, 88(4), 257–263. doi:88/4/257.1..

McCauley, D. E. (1994). Contrasting the distribution of chloroplast DNA and allozyme polymorphism among local populations of Silene alba: implications for studies of gene flow in plants. Proceedings of the National Academy of Sciences, 91(17), 8127–8131.

McCauley, D. E., Raveill, J., & Antonovics, J. (1995). Local founding events as determinants of genetic structure in a plant metapopulation. Heredity, 75(6), 630–636. doi:doi:10.1038/hdy.1995.18.

Mitchell, C. E., Agrawal, A. A., Bever, J. D., Gilbert, G. S., Hufbauer, R. A., Klironomos, J. N., … Vázquez, D. P. (2006). Biotic interactions and plant invasions. Ecology Letters, 9(6), 726–740. doi:10.1111/j.1461-0248.2006.00908..

Murren, C. J., & Dudash, M. R. (2012). Variation in inbreeding depression and plasticity across native and non-native field environments. Annals of Botany, 109(3), 621–632. doi:10.1093/aob/mcr32.

Normand S., Zimmermann N. E., Schurr F. M., & Lischke H. (2014). Demography as the basis for understanding and predicting range dynamics. Ecography, 37(12), 1149–1154. doi:10.1111/ecog.0149.

Orians, C. M., & Ward, D. (2010). Evolution of plant defenses in nonindigenous environments. Annual Review of Entomology, 55(1), 439–459. doi:10.1146/annurevento-112408-08533.

Orr, H. A. (2009). Fitness and its role in evolutionary genetics. Nature Reviews Genetics, 10(8), 531–539. doi:10.1038/nrg260.

Phillips, B. L., Brown, G. P., & Shine, R. (2010). Life-history evolution in range-shifting populations. Ecology, 91(6), 1617–1627. doi:10.1890/09-0910..

R Development Core Team. (2015). R: a language and environment for statistical computing. R Foundation for Statistical Computing. Retrieved from https://www.R-project.org/

Reed, D. H., Briscoe, D. A., & Frankham, R. (2002). Inbreeding and extinction: the effect of environmental stress and lineage. Conservation Genetics, 3(3), 301–307. doi:10.1023/A:101994813026.

Reed, D. H., Fox, C. W., Enders, L. S., & Kristensen, T. N. (2012). Inbreeding–stress interactions: evolutionary and conservation consequences. Annals of the New York Academy of Sciences, 1256(1), 33–48. doi:10.1111/j.1749-6632.2012.06548..

Richards, C. M. (2000). Inbreeding depression and genetic rescue in a plant metapopulation. The American Naturalist, 155(3), 383–394. doi:10.1086/30332.

Rius, M., & Darling, J. A. (2014). How important is intraspecific genetic admixture to the success of colonising populations? Trends in Ecology & Evolution, 29(4), 233–242. doi:10.1016/j.tree.2014.02.00.

Roman, J., & Darling, J. (2007). Paradox lost: genetic diversity and the success of aquatic invasions. Trends in Ecology & Evolution, 22(9), 454–464. doi:10.1016/j.tree.2007.07.00.

Rosche, C., Hensen, I., & Lachmuth, S. (2017). Local pre-adaptation to disturbance and inbreeding–environment interactions affect colonisation abilities of diploid and tetraploid Centaurea stoebe. Plant Biology. doi:10.1111/plb.1262.

Sandner, T. M., & Matthies, D. (2016). The effects of stress intensity and stress type on inbreeding depression in Silene vulgaris. Evolution, 70(6), 1225–1238. doi:10.1111/evo.1292.

Sandner, T. M., & Matthies, D. (2017). Interactions of inbreeding and stress by poor host quality in a root hemiparasite. Annals of Botany, 119(1), 143–150. doi:10.1093/aob/mcw19.

Schou, M. F., Loeschcke, V., & Kristensen, T. N. (2015). Inbreeding depression across a nutritional stress continuum. Heredity, 115(1), 56–62. doi:10.1038/hdy.2015.1.

Schrieber, K., Kröner, L., Schweiger, R. & Müller, C. (accepted for publication in Journal of Ecology): Inbreeding diminishes herbivore-induced metabolic responses in native and invasive plant populations.

Schrieber, K., & Lachmuth, S. (2017). The Genetic Paradox of Invasions revisited: the potential role of inbreeding × environment interactions in invasion success. Biological Reviews, 92(2), 939–952. doi:10.1111/brv.1226.

Schrieber, K., Wolf, S., Wypior, C., Höhlig, D., Hensen, I., & Lachmuth, S. (2017). Adaptive and non-adaptive evolution of trait means and genetic trait correlations for herbivory resistance and performance in an invasive plant. Oikos, 126(4), 572–582. doi:10.1111/oik.0378.

Schultz, E. L., Eckberg, J. O., Berg, S. S., Louda, S. M., & Miller, T. E. X. (2017). Native insect herbivory overwhelms context dependence to limit complex invasion dynamics of exotic weeds. Ecology Letters, 20(11), 1374–1384. doi:10.1111/ele.1283.

Taylor, D. R., & Keller, S. R. (2007). Historical range expansion determines the phylogenetic diversity introduced during contemporary species invasion. Evolution, 61(2), 334–345. doi:10.1111/j.1558-5646.2007.00037..

Travis, J. M. J., Münkemüller, T., Burton, O. J., Best, A., Dytham, C., & Johst, K. (2007). Deleterious mutations can surf to high densities on the wave front of an expanding population. Molecular Biology and Evolution, 24(10), 2334–2343. doi:10.1093/molbev/msm16.

Venables, W. N., & Ripley, B. D. (2000). Modern Applied Statistics with S. Fourth Edition. New York, USA: Springer.

Whitney, K. D., & Gabler, C. A. (2008). Rapid evolution in introduced species, ‘invasive traits’ and recipient communities: challenges for predicting invasive potential. Diversity and Distributions, 14(4), 569–580. doi:10.1111/j.1472-4642.2008.00473..

Wolfe, L. M. (2002). Why alien invaders succeed: support for the escape-from-enemy hypothesis. The American Naturalist, 160(6), 705–711. doi:10.1086/34387.

Wolfe, L. M., Elzinga, J. A., & Biere, A. (2004). Increased susceptibility to enemies following introduction in the invasive plant Silene latifolia. Ecology Letters, 7(9), 813–820. doi:10.1111/j.1461-0248.2004.00649..

Zhang, D.-Y., & Jiang, X.-H. (2006). Interactive effects of habitat productivity and herbivore pressure on the evolution of anti-herbivore defense in invasive plant populations. Journal of Theoretical Biology, 242(4), 935–940. doi:10.1016/j.jtbi.2006.05.01.

Zuur, R. A. F., Ieno, E. N., Walker, N. J., Saviliev, A. A., & Smith, G. M. (2009). Mixed Effects Models and Extensions in Ecology. New York: Springer.

